# The impact of 10-valent Pneumococcal Conjugate Vaccine on the incidence of radiologically-confirmed pneumonia and clinically-defined pneumonia in Kenyan children

**DOI:** 10.1101/369686

**Authors:** Micah Silaba, Michael Ooko, Christian Bottomley, Joyce Sande, Rachel Benamore, Kate Park, James Ignas, Kathryn Maitland, Neema Mturi, Anne Makumi, Mark Otiende, Stanley Kagwanja, Sylvester Safari, Victor Ochola, Tahreni Bwanaali, Evasius Bauni, Fergus Gleeson, Maria Deloria Knoll, Ifedayo Adetifa, Kevin Marsh, Thomas N Williams, Tatu Kamau, Shahnaaz Sharif, Orin S Levine, Laura L Hammitt, J Anthony G Scott

## Abstract

**Background:** Pneumococcal conjugate vaccines (PCV) are highly protective against invasive pneumococcal disease caused by vaccine serotypes but the burden of pneumococcal disease in developing countries is dominated by pneumonia, most of which is non-bacteraemic. We examined the impact of PCV on pneumonia incidence.

**Methods:** We linked prospective hospital surveillance for clinically-defined WHO severe or very-severe pneumonia at Kilifi County Hospital from 2002-2015 to population surveillance at Kilifi Health and Demographic Surveillance System, comprising 45,000 children aged <5 years. Chest radiographs were read according to a WHO standard. A 10-valent pneumococcal non-typeable *Haemophilus influenzae* protein D conjugate vaccine (PCV10) was introduced in Kenya in January 2011. In Kilifi, there was a catch-up campaign for children aged <5 years. We estimated the impact of PCV10 on pneumonia incidence through interrupted time series analysis accounting for seasonal and temporal trends.

**Findings:** The incidence of admission with clinically-defined pneumonia in 2002/3 was 21·7/1000/year in children aged 2-59 months. This declined progressively over 13 years. By the end of March 2011, 61·1% of children aged 2-11 months received ≥2 doses and 62·3% of children aged 12-59 months received ≥1 dose of PCV10. Adjusted incidence rate ratios for admissions with radiologically-confirmed pneumonia, clinically-defined pneumonia, and diarrhoea (control condition), associated with PCV10 introduction, were 0·52 (95% CI 0·32-0·86), 0·73 (95% CI 0·54-0·97) and 0·63, (95% CI 0·31-1·26), respectively. The annual incidence of clinically-defined pneumonia in December 2010 was 12·2/1000; this was reduced by 3·3/1000 with PCV10 introduction.

**Interpretation:** Over 13 years, hospitalisations for clinically-defined pneumonia declined progressively at Kilifi County Hospital but fell abruptly by 27% in association with PCV10 introduction. The incidence of radiologically-confirmed pneumonia fell by 48%. The burden of childhood pneumonia in Kilifi, Kenya, has been reduced substantially by PCV10.

**Funding:** Gavi, Wellcome Trust

## Introduction

Pneumonia is the single greatest cause of death in children^1^ and the commonest cause of fatal pneumonia is *Streptococcus pneumoniae*^2^.

Pneumococcal Conjugate Vaccines (PCV) are highly efficacious against invasive pneumococcal disease (IPD) caused by vaccine serotypes. The introduction of PCVs in high-income countries has decreased transmission of vaccine serotypes and reduced IPD among both vaccinated and unvaccinated populations. However, IPD represents only a small fraction the burden of pneumococcal disease. In a randomised-controlled trial (RCT) of 9-valent PCV in The Gambia, 15 cases of radiologically-confirmed pneumonia were prevented for every 2 cases of IPD^3^.

The efficacy of PCV against clinically-defined pneumonia is low (0-17%) in RCTs in developing countries^3–5^. This suggests that clinically-defined pneumonia as an endpoint has poor specificity for pneumococcal pneumonia. Recognising this limitation, the World Health Organization (WHO) developed a set of interpretive criteria and procedures to standardize the reading of paediatric chest radiographs in pneumonia cases^6, 7^ which defined an endpoint that has higher specificity for pneumococcal pneumonia and commensurately higher estimates of vaccine efficacy (20-37%)^3, 4, 8, 9^.

Longitudinal before-after studies of disease incidence, with an interrupted time series analysis are likely to capture the beneficial effects of herd protection due to reduced transmission of vaccine-serotype pneumococci as well as the effects of direct vaccine protection. They are also sensitive to serotype replacement diseasse if infection with non-vaccine serotypes leads to pneumonia. To date there have been only two field studies of PCV effectiveness against pneumonia in low-income settings; one had just two years of pre-vaccine surveillance, the other none^10, 11^.

In America, the impact of PCV7 on routine hospital admissions with all-cause pneumonia was estimated, using interrupted time-series analysis, as a 39% reduction in children aged <2 years which is a substantially greater than the efficacy estimates against clinical pneumonia or radiologically-confirmed pneumonia in an RCT^12–14^. Unfortunately, in most low-income settings, the quality of routine administrative hospital data is insufficient for evaluation using this study design.

In interrupted time-series analyses, after adjusting for seasonal and temporal trends in pneumonia hospitalization, the residual change in incidence associated with vaccine introduction is as robust an estimate of vaccine impact as is possible in a non-randomised design^15, 16^. We used this method to capture the impact of PCV10 on both clinically-defined and radiologically-confirmed pneumonia in a unique clinical and demographic surveillance platform with real-time monitoring of vaccine coverage^17, 18^. We introduced PCV10 with a catch-up campaign for children aged <5 years to increase temporal specificity of the time series effect.

## Methods

### Setting

We studied residents of the Kilifi Health and Demographic Surveillance System (KHDSS) aged ≥2 months to <12 years. KHDSS has monitored births, deaths and migration events in a population of 280,000 through 4-monthly household visits since 2001^18^.

Kilifi County Hospital (KCH) is centrally located within KHDSS and is the only paediatric inpatient facility in the study area. It has 55 paediatric beds. Since 2002 all admissions have been recorded using a standard electronic clinical record linked to the KHDSS population register.

We used World Health Organization (WHO) definitions of clinical pneumonia applicable at the start of the surveillance based on presentation with cough or difficulty breathing^19^. Those with lower chest-wall indrawing but no danger signs had an admission diagnosis of severe pneumonia; those with ≥1 danger sign had very severe pneumonia. The danger signs were central cyanosis, inability to drink, convulsions, lethargy, prostration, unconsciousness or head nodding^19^. As there were no specific definitions for children aged 5-11 years, we applied the same criteria as for younger children. Children with non-severe pneumonia are not normally admitted and were not included in the surveillance. Hospitalisation with diarrhoea served as a control condition. Diarrhoea was defined as ≥3 loose stools in the last 24 hours.

From April 2006 onwards, children with WHO-defined severe or very-severe pneumonia were investigated, whenever possible, with a single anterior-posterior chest radiograph. Presentation with convulsions alone, or lethargy alone, without other signs of pneumonia, was not considered by the local ethical review committee to be sufficient justification for investigation with a chest radiograph.

Chest radiographs were taken with a Philips Cosmos-BS machine or a Philips Practix 360 portable machine, which became available in March 2012. The radiology system was digitised in August 2011 and thereafter a Philips PCR Eleva-S was used to process digital cassettes (10 x 12 inches; 1670 x 2010 pixels). Archived film images were digitised using a Vidar Pro Advantage digitizer. Images were encoded using Hipax software into DICOM format at 150dpi and 12bits. All images were cropped to de-identify the patient and remove peripheral clues about the radiological method used, then distributed in JPEG format in batches of 100 selected at random from pre- and post-vaccine introduction images.

### Radiological reading and interpretation

Radiological interpretation followed the standard defined by WHO for identification of primary end-point pneumonia (PEP) which was used in the phase III trials of pneumococcal conjugate vaccines^6, 7^. All images were first categorized by quality; adequate/optimal, sub-optimal or unreadable/uninterpretable. Uninterpretable images were not assigned a diagnosis. PEP was defined by the presence of consolidation and/or pleural effusion.

Each image was read independently by two primary readers, a consultant radiologist and a trainee paediatrician in Kenya. All images with discordant interpretations, and 10-15% of those with agreement, were referred to three consultant radiologists in Oxford who arbitrated the readings. Concordant readings of the primary readers were considered final.

### Vaccine introduction and monitoring

In January 2011, a 10-valent PCV (Synflorix®), consisting of capsular polysaccharides of serotypes 1, 4, 5, 6B, 7F, 9V, 14, 18C, 19F and 23F conjugated to either non-typeable *Haemophilus influenzae* (NTHi) protein D, diphtheria or tetanus toxoid, was introduced in Kenya in three doses at 6, 10 and 14 weeks; there was a 3-dose catch-up campaign for infants during 2011. In addition, in Kilifi County, children aged 12-59 months were offered two doses of vaccine through mass-campaigns on 31^st^ January-6^th^ February 2011 and 21^st^-27^th^ March 2011.

*Haemophilus influenzae* type b conjugate vaccine was introduced in Kenya in 2001. Rotavirus vaccine (Rotarix®) was introduced, without a catch-up campaign, in July 2014.

Vaccine surveillance was established in April 2009 in 26 vaccine clinics serving KDHSS^17^. Data clerks recorded all immunisations given against the identity of the child in the KHDSS population register at the point of vaccination. However, during the catch-up campaign, vaccinations were recorded against lists of KHDSS residents.

### Statistical analysis

The incidence of hospitalization with clinically-defined pneumonia or diarrhoea was calculated for each month between May 2002 and March 2015. Children who had both pneumonia and diarrhoea were classified as having pneumonia alone. The monthly incidence of radiologically-confirmed pneumonia was calculated between April 2006 and March 2014. Mid-month population counts from the KHDSS were used to estimate child years at risk in each month.

Linear regression models were fitted to log-transformed monthly rates of radiologically-confirmed and clinically-defined pneumonia to estimate the effect of PCV10. The models included a period effect (pre- and post-PCV10), monthly time trend and seasonality, which was modelled using month of year. Differences in the time trend before and after vaccination were tested as an interaction. We modelled the error as an autoregressive moving average (ARMA) process, using Aikake’s information criterion and plots of the autocorrelation function of residuals to choose the order of the process^20^.

We excluded January to March 2011 as a transition period during which PCV10 was introduced among children under 5 years. We also excluded admissions in December 2012 and December 2013 because of nurses’ strikes.

Where radiographs were not obtained, we used multiple imputations based on information on admission year, month of admission, sex, age, HIV status, malaria slide positivity, pneumonia severity, outcome of hospitalization (alive at discharge or died), referral, and day of the week when the patient was admitted. Twenty imputed datasets were created, using the method of chained equations^21, 22^, and Rubin’s rules were used to combine estimates across the imputed datasets.

Statistical analyses were done in STATA (v14).

Written informed consent was obtained from the parents/guardians of all participants in the study. The study was approved by the KEMRI National Ethical Review Committee and Oxford Tropical Research Ethics Committee.

The study was funded by Gavi, The Vaccine Alliance and The Wellcome Trust. The corresponding author had full access to all the data in the study and had final responsibility for the decision to submit for publication.

## Results

In May 2002, there were 37,556 residents in Kilifi HDSS aged 2-59 months and 44,672 residents aged 60-143 months. By March 2015 these figures were 45,601 and 62,502, respectively (Table 1). By the end of March 2011, 61·1% of children aged 2-11 months received at least two doses of PCV10 and 62·3% of children aged 12-59 months received at least one dose of PCV10. Subsequently, coverage increased slowly throughout the remainder of the study period (Table 1).

**Table 1.**
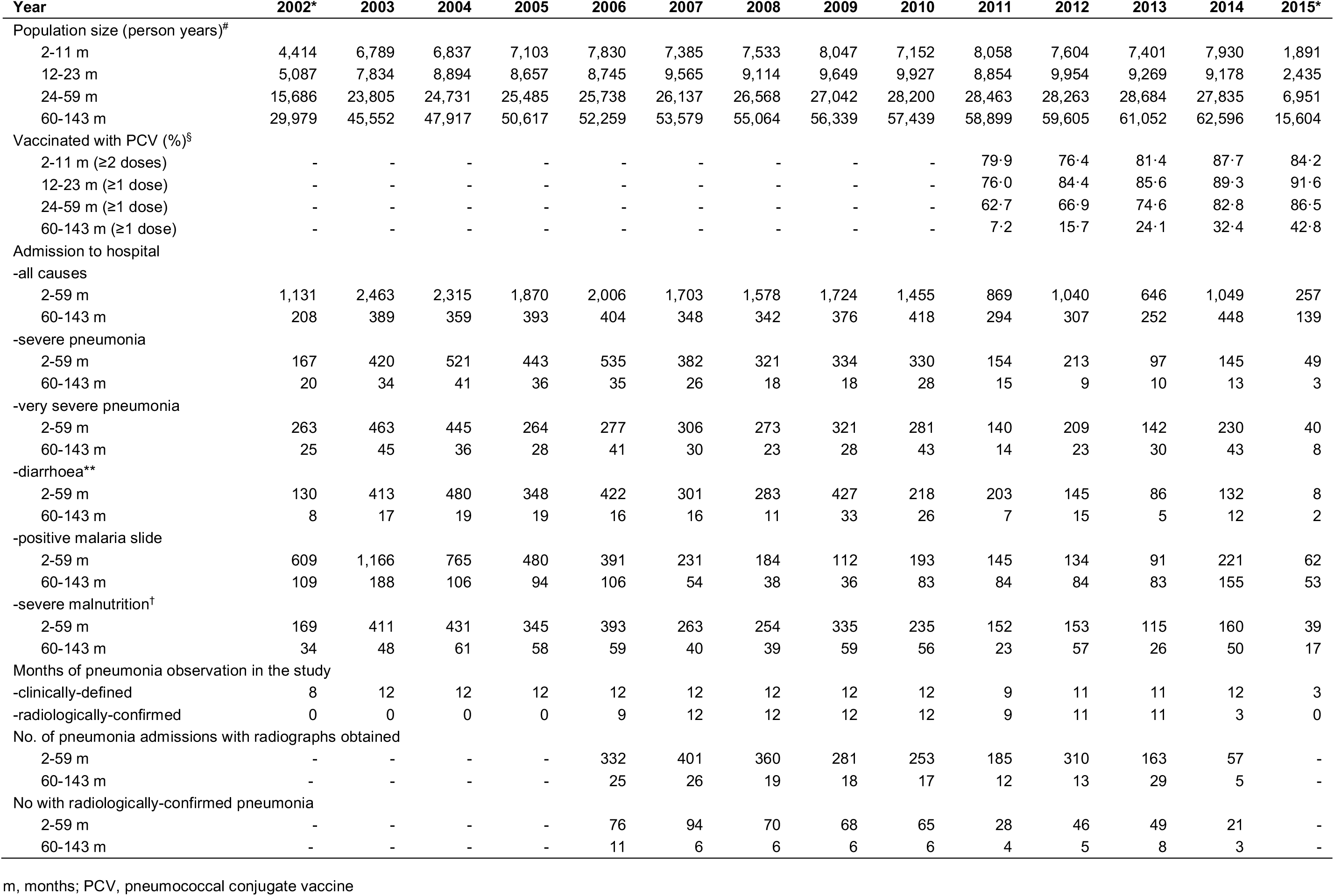
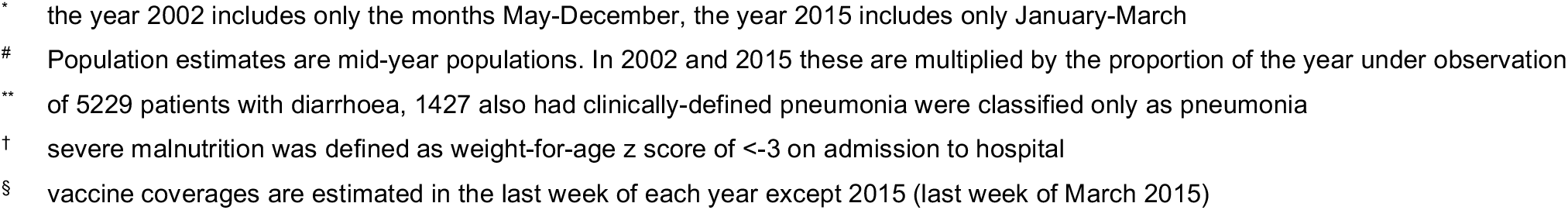
Population size under observation, coverage (%) of PCV10 and numbers of admissions to hospital by year of surveillance, among children resident within the Kilifi Health and Demographic Surveillance System.

Between May 2002 and March 2015, inclusive, 44,771 children aged 2-143 months were admitted to Kilifi County Hospital (Table 1). We excluded 810 admissions between January and March 2011 and 182 admissions during nurses’ strikes. Of the remaining 43,779 admissions, 24,783 (57%) were residents of KHDSS (Figure 1) and, of these, 8,488 (34%) had severe or very severe clinical pneumonia. Throughout the 13-year study period the numbers of admissions to hospital among KHDSS residents aged 2-59 months fell progressively, particularly those with severe malnutrition (defined as weight for age z-score <-3) and a positive malaria slide (Table 1). The prevalence of HIV infection among mothers attending the ante-natal clinic in Kilifi County Hospital was 4·1% in 2005-7, 4·2% in 2008-10 and 2·7% in 2011-15.

**Figure 1.**
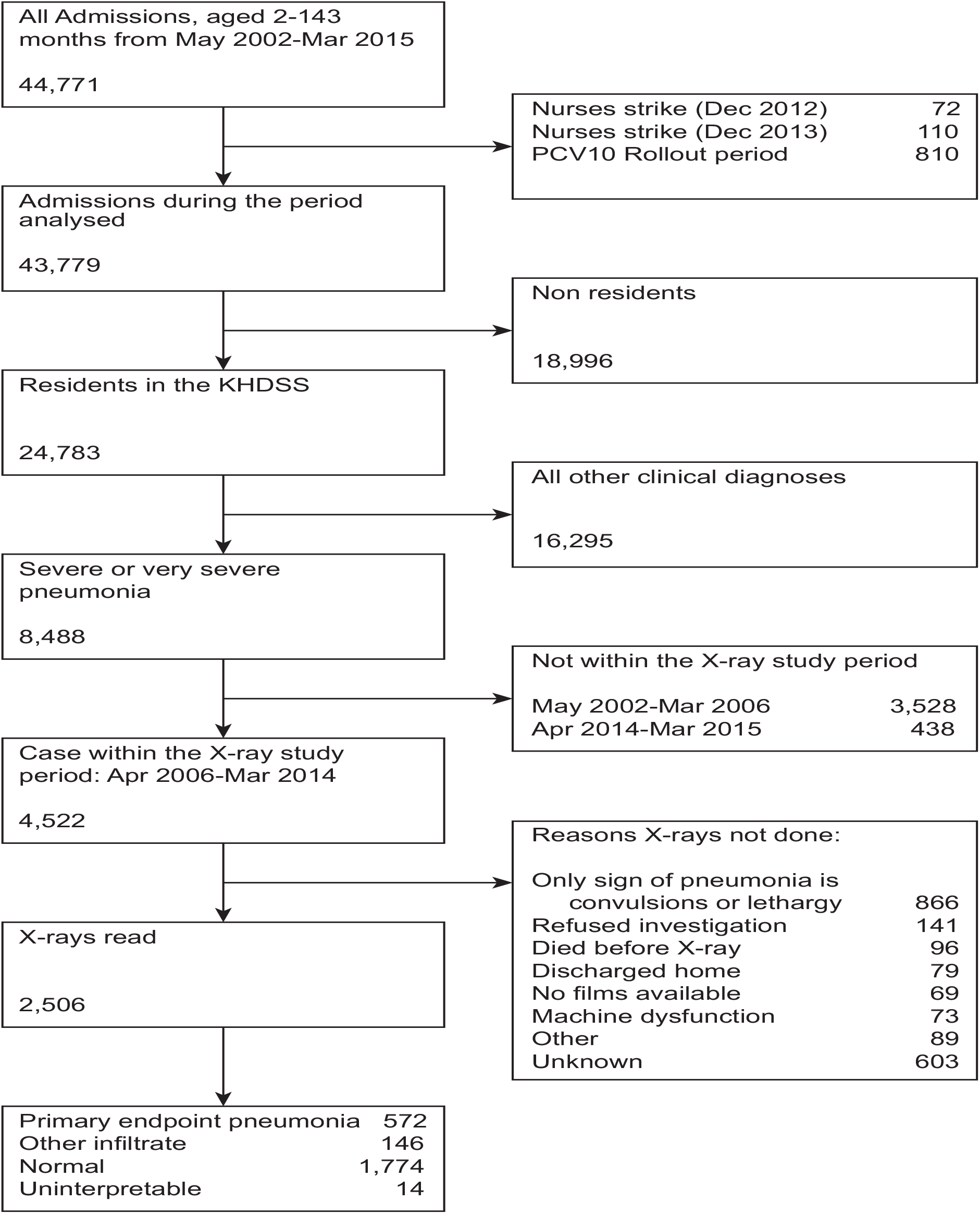
Flow chart illustrating the selection of patients for radiographs and the reasons for missing radiographs.

Among pneumonia patients, 4,522 (53%) were admitted during the radiological study period (April 2006-March 2014) and ≥1 chest radiograph was obtained from 2,506 (55·4%) overall; radiographs were obtained from 51% (1732/3373) and 67% (774/1149) in the pre-vaccine and post-vaccine periods, respectively. Among the 2,506 radiographs read, 14 were designated as unreadable or uninterpretable, 572 had primary end-point pneumonia (PEP), 146 had other infiltrates and 1,774 were normal. Prior to 2011, primary end-point pneumonia was identified in 21·3% (185/867), 25·8% (109/423), 24·2% (79/327) and 24·0% (25/104) of readable radiograph images among children aged 2-11m, 12-23m, 24-59m and 60-143m, respectively.

Prior to imputation and modelling, the crude incidence rates for radiologically-confirmed pneumonia among children aged 2-59 months were 180·9 (95% CI 163·0-200·2) and 110·7 (95% CI 93·3-130·3) per 100,000 person years for the periods before and after PCV10 introduction, respectively; the crude Incidence Rate Ratio (IRR) was 0·61 (95% CI 0·50-0·74). In the interrupted time-series model, which adjusted for season and time (Figure 2, Table 2) the IRR for radiologically-confirmed pneumonia associated with PCV10 introduction was 0·52 (95% CI 0·32-0·86). The impact was greatest among those aged 12-59 months and there was no evidence of impact among children aged 5-11 years (Table 2). After accounting for season and the vaccine effect, the underlying incidence of radiologically-confirmed pneumonia among children 2-59 months was stable over time (IRR per year 0·98, 95% CI 0·88- 1·09).

**Figure 2.**
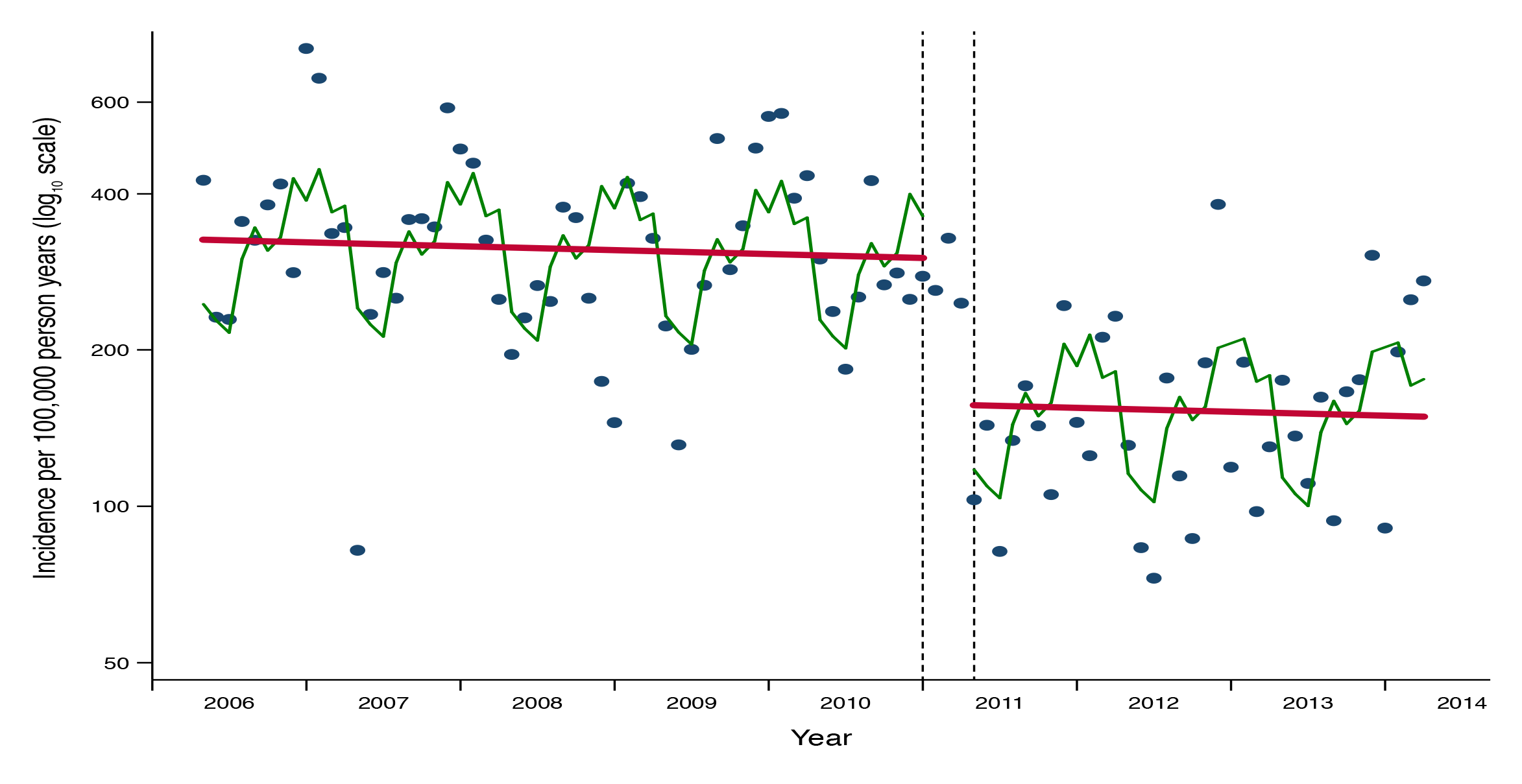
Monthly incidence of admission to Kilifi County Hospital among children aged 2-59 months with WHO-defined radiologically-confirmed pneumonia and modelled predictions. Blue dots are monthly incidence rates, the green line is the regression line for season and the red line is the regression line for temporal trend. The black dashed lines define the transition period during which PCV10 was introduced among children under 5 years.

**Figure 3.**
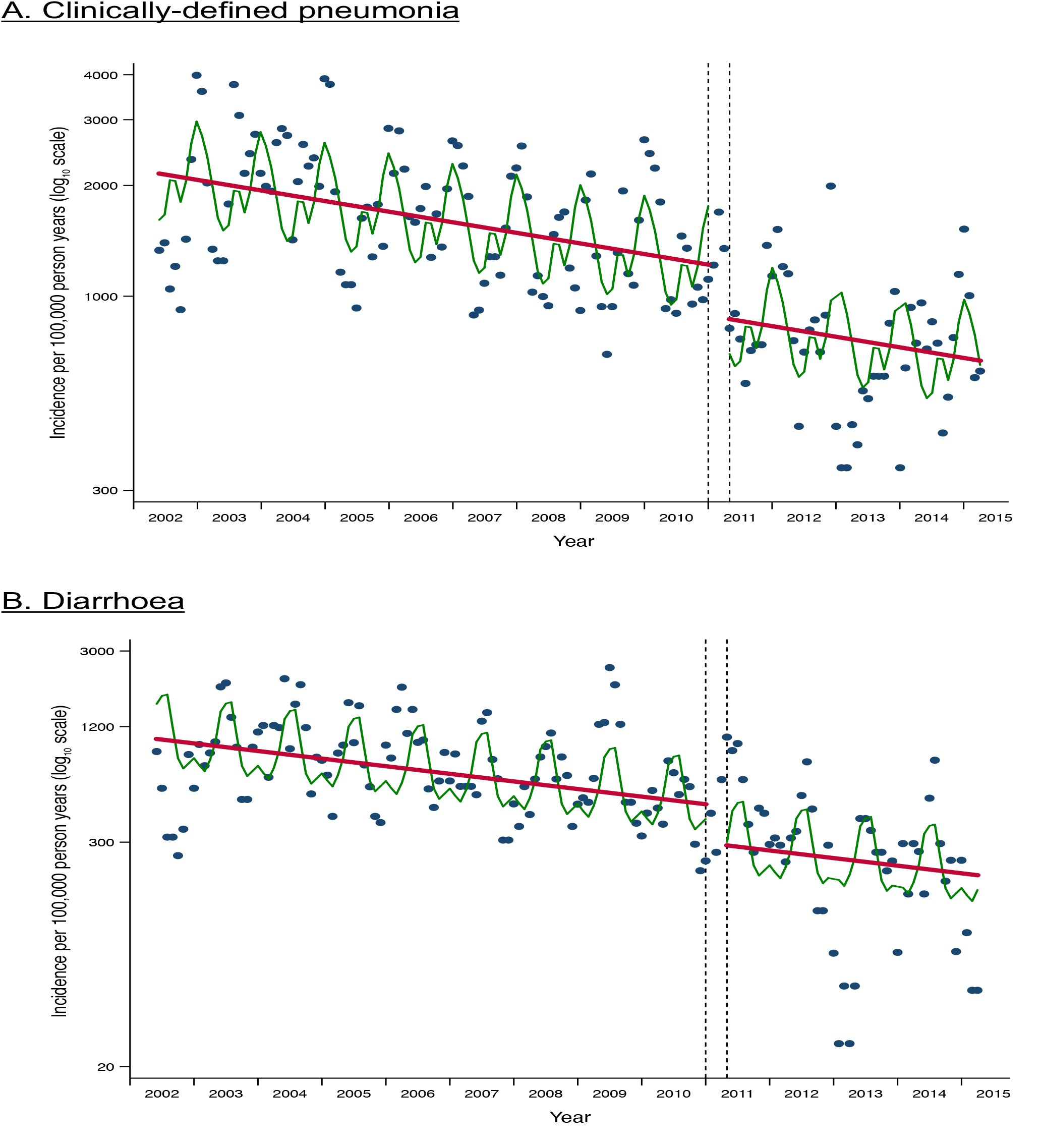
Monthly incidence of admission to Kilifi County Hospital among children aged 2-59 months with A. Clinically-defined pneumonia (WHO severe or very severe pneumonia) and B. Diarrhoea. Blue dots are monthly incidence rates, the green line is the regression line for season and the red line is the regression line for temporal trend. The black dashed lines define the transition period during which PCV10 was introduced among children under 5 years.

**Table 2.** Incidence rate ratios for the effects of PCV10 introduction against hospitalisation with radiologically-confirmed pneumonia among children aged 2-143 months in strata of age.

| Sub-groups* | Incidence Rate Ratio for PCV10 introduction | 95% CI | P value |
| --- | --- | --- | --- |
| Age |  |  |  |
| 2-59 months (primary analysis) | 0·52 | 0·32, 0·86 | 0·011 |
| 2-11 months | 0·73 | 0·39, 1·36 | 0·324 |
| 12-23 months | 0·54 | 0·27, 1·10 | 0·091 |
| 24-59 months | 0·50 | 0·24, 1·01 | 0·052 |
| 60-143 months | 0·89 | 0·47, 1·69 | 0·724 |
| Clinical severity (2-59 months only) |  |  |  |
| Severe pneumonia | 0·56 | 0·30, 1·06 | 0·077 |
| Very severe pneumonia | 0·52 | 0·28, 0·96 | 0·038 |

\*Interrupted time series analysis adjusted for season (month of year) and study time (months).

The annual incidence of admission with clinically-defined pneumonia in 2002/3 was 2,170/100,000 in children aged 2-59 months and there was a significant reduction in incidence of admission to hospital across the study period of 1% per month; pneumonia admissions were also subject to marked seasonal variation with the highest rates in November to January and the lowest in April to June (Suppl table 1). After adjusting for these factors, the IRR for admissions with severe or very severe pneumonia associated with introduction of the PCV10 programme was 0·73 (95%CI 0·54-0·97). Vaccine impact was greater for severe pneumonia than for very severe pneumonia (Table 3). There was no evidence of benefit to children aged 5-11 years.

**Table 3.**
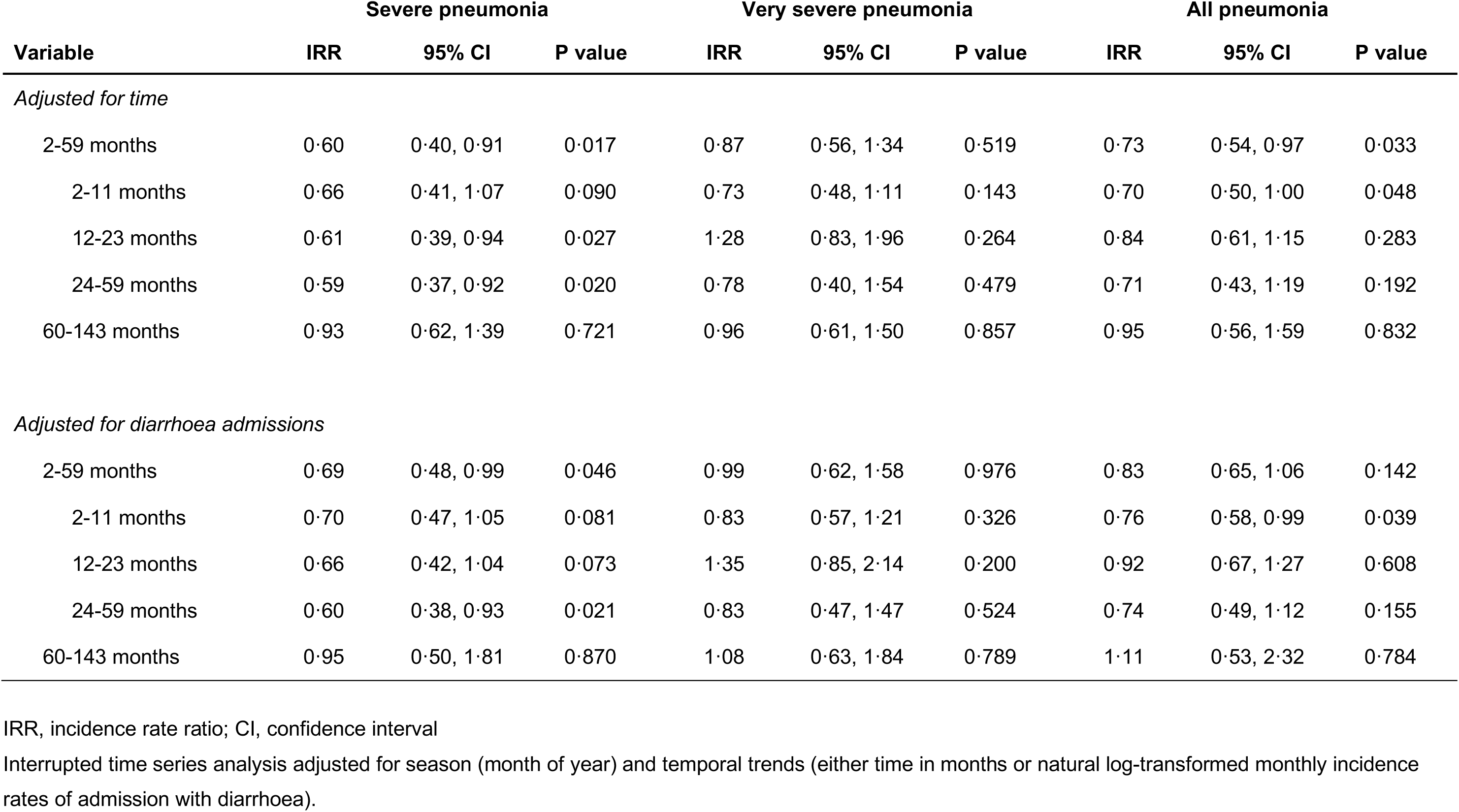
Incidence rate ratios for the effects of PCV10 introduction against hospitalisation with WHO-defined severe or very severe pneumonia among children aged 2-143 months.

The control condition, incidence of admission with diarrhoea among children aged 2- 59 months, was not associated with PCV10 introduction (IRR 0·63, 95% CI 0·31- 1·26). However, in age-stratified analyses, there was a significant association between PCV10 introduction and incidence of diarrhoeal admissions in those aged 12-23 months, 24-59 months and 5-11 years (Suppl table 3). The IRR among infants (0·79) was considerably less extreme than the IRRs observed in older age groups (0·62-0·66). Truncating analysis time at the point of Rotavirus Vaccine introduction did not significantly alter these findings.

There was no significant interaction between study time and vaccine era in the analysis of the incidence of clinically-defined pneumonia nor of radiologically-confirmed pneumonia. This suggests there was little further development of indirect protection after the catch-up campaign.

The absolute reduction in admission rates was estimated by applying the vaccine effectiveness estimates to the model predictions of incidence in the month immediately prior to PCV10 introduction (December 2010). The modelled incidence rates of clinically-defined and radiologically-confirmed pneumonia were 1,220 and 301 per 100,000 person years, respectively, among children aged <5 years. The vaccine preventable disease incidence was 329 and 144 cases per 100,000 person years, respectively.

We explored whether the observed impact of PCV10 on diarrhoea in some age groups implied residual confounding in hospital presentation patterns. To select a suitable control condition, we examined the correlation between annual counts of admissions with clinically-defined pneumonia in the pre-vaccine period against annual counts of admissions with other discharge diagnoses. The greatest correlations were found with unclassified discharges and with gastroenteritis, though unclassified discharges were relatively uncommon. The correlation with the admission diagnosis diarrhoea was greater yet (Suppl Table 2). After adjusting for log-transformed monthly rates of diarrhoea admissions, instead of time in months, the IRR for severe pneumonia associated with PCV10 in children aged 2-59 months was 0·69 (95% CI 0·48-0·99); there was no observed effect of PCV10 on the incidence of very severe pneumonia nor on either grade of pneumonia in children aged 5-11 years (Table 3).

## Discussion

Analysis of the rates of hospital admission from a rolling cohort of approximately 43,000 children aged 2-59 months in Kilifi, Kenya, suggests that the introduction of PCV10, with a simultaneous catch-up campaign for children under 5 years, was associated with a reduction in childhood hospitalizations with clinically-defined and radiologically-confirmed pneumonia of 27% and 48%, respectively. The vaccine reduced the incidence of admission to hospital with clinically-defined and radiologically-confirmed pneumonia by 329 and 144 per 100,000 person years, respectively. There was no evidence of an impact among children aged ≥5 years.

The impact observed in Kilifi was considerably greater than the vaccine efficacy estimates from individually randomized controlled trials (RCTs) of PCVs. Against severe clinical pneumonia, the vaccine efficacy of a 9-valent PCV was 12% (95% CI-9, 29) in The Gambia and 17% (95% CI 7, 26) in South Africa^3, 5^. Against radiologically-confirmed pneumonia the vaccine efficacies in The Gambia and South Africa were 37% and 20%, respectively^3, 8^. In Bohol, The Philippines, the point estimate for vaccine efficacy of an 11-valent PCV against radiologically-confirmed pneumonia was 22·9% (95% CI-1·1, 41·2); there was no protection against the clinically-defined pneumonia^4^. A RCT of PCV10, conducted in Argentina, Panama and Columbia estimated vaccine efficacy against radiologically-confirmed pneumonia at 22·4%^9^. These vaccine efficacy estimates, derived from individually-randomised trials, measure only the direct protective effect of the vaccine whereas the impact estimates in this report combine both direct and indirect effects. In America, the indirect effect of PCV7 against pneumonia was substantial^23^ and it is not unexpected, therefore, that the impact estimates in a real-world implementation are considerably greater than the efficacy estimates of the RCTs.

There is relatively little information on PCV impact elsewhere in tropical Africa. In a retrospective analysis (2002-2012) of clinical pneumonia among admission case-records from 5 district hospitals in Rwanda, the impact of PCV7, introduced in 2009, was similar to that observed in Kilifi (Vaccine effectiveness 54%, 95% CI 42, 63%)^24^. However, an impact study of PCV in children aged <5 years in The Gambia, comparing rates in the post-PCV13 era (2014-2015) with those in the pre-PCV7 era (2008-2010), found only a 5-15% reduction in hospitalised clinical pneumonia, depending on age. For children hospitalised with radiologically-confirmed pneumonia the reductions were 24-31%^10^, which were lower than the 37% estimate derived from an RCT of PCV9 in the same setting^3^.

The PCV10 introduced in Kenya uses protein D of *Haemophilus influenzae* as a carrier protein for 8 of the 10 polysaccharide antigens. This protein is immunologically exposed on the bacterial surface and antibodies to protein D may protect against infections of unencapsulated (non-typeable) *H. influenzae*. However, *H. influenzae* is rarely identified by blood culture as a cause of pneumonia in Kilifi^26^ and the contribution of protein D to the impact on radiologically-confirmed pneumonia in this study is likely to be small.

In Kilifi, across the whole 13-year study period, there was a sustained reduction in admission incidence rates, particularly for malaria and malnutrition, but also for clinically-defined pneumonia. Over the same period infant and under five mortality ratios declined by more than 50%^27^ arguing that changing admission rates reflect a genuine improvement in health rather than a change in health-seeking behaviour. In fact, the impact of general health trends on the incidence of clinically-defined pneumonia in the pre-vaccine period was greater than the impact of PCV10 at introduction in 2010. Among children 2-59 months old, the annual incidence of clinically-defined pneumonia per 100,000 children declined from 2,170 to 1,200 between 2002/3 and in December 2010; with the introduction of PCV10 it declined by a further 329. In other settings, which have a higher baseline incidence, the absolute benefits of PCV10 are likely to be considerably greater. However, the full magnitude of the impact may take longer in the absence of a catch-up campaign.

The observed impact of PCV10 against radiologically-confirmed pneumonia is substantially greater than that against clinically-defined pneumonia suggesting that vaccine serotype pneumococci account for proportionately more cases of radiologically-confirmed pneumonia than of clinically-defined pneumonia. The WHO radiological standard was developed to generate an endpoint that was specific for bacterial pneumonia and, in the presence of a vaccine programme for *H. influenzae* type b, it is relatively specific for pneumococcal pneumonia^3, 6^. However, as well as differences in impact, we observed marked differences in temporal trends; radiologically-confirmed pneumonia was stable whilst clinically-defined pneumonia declined sharply with time. The clinical presentations of pneumonia and malaria are difficult to distinguish^28^ and the prevalence of malaria has declined sharply from 1999-2007^29^ suggesting that some of the temporal trend in clinically-defined pneumonia admissions may be attributable to changes in malaria incidence.

Following the example of previous studies^12^ we selected diarrhoea as a control condition because it was common and should be unaffected by PCV10. However, we observed an unexpected fall in diarrhoea admissions among children 2-59 months associated with the timing of PCV10 introduction (IRR 0·63 95% CI 0·31, 1·26). We examined whether there was residual confounding in the temporal pattern of hospital presentations, by adjusting for diarrhoea admissions, instead of time in years. The impact against severe pneumonia was reduced slightly from IRR 0·60 to 0·69 but the impact on very severe pneumonia disappeared.

The decline in diarrhoeal admissions following PCV10 introduction is unlikely to be due to a biological effect of PCV10. However, it may be a marker of another intervention in this area, targeting diarrhoea, contemporaneous with PCV10 introduction. The effect was significant in all age groups beyond infancy and especially marked in those aged 5-11 years, who were too old to be vaccinated with PCV10 (Suppl Table 3). We were not aware of any health intervention targeting diarrhoea at this time, but if there was, then selecting diarrhoea as an adjustment variable for temporal trends in hospital presentation would act to underestimate the true impact of PCV10 against clinically-defined pneumonia. Despite the ambiguity of the diarrhoea results, there is no question that the vaccine had a large and significant impact against severe pneumonia in children aged <5 years. In addition, the impact against radiologically-confirmed pneumonia requires no adjustment as there was no temporal trend in disease incidence (Figure 2).

Several features of the present study argue strongly that the associations observed between vaccine introduction and disease incidence were causal: the duration of surveillance was long and the surveillance methods were consistent; the vaccine programme was introduced with a rapid catch-up campaign and nearly two thirds of the target population were given an immunizing schedule of vaccine at the same time point; the analyses took careful account of long-term trends in disease incidence and seasonal variation; the magnitude of the effects was very large and therefore difficult to attribute to incidental unobserved improvements in the environment. In addition, these changes in pneumonia incidence occurred simultaneously with an abrupt 64% reduction in the prevalence of carriage of the serotypes included in the vaccine^25^ and with a 68% decline in the frequency of all IPD^30^.

Taken together these factors suggest that the introduction of PCV10 in Kilifi, Kenya has reduced the incidence of hospitalizations with clinically-defined pneumonia by approximately one quarter, and of radiologically-confirmed pneumonia by approximately half. Given that pneumonia, rather than IPD, is where the greatest burden of pneumococcal disease lies, these findings imply a very considerable improvement in child health associated with the implementation of a PCV10 programme.

## Acknowledgements

We would like to thank all the clinical and field staff involved in collecting surveillance data and the patients and their families for their cooperation with the surveillance in Kilifi. This paper is published with the approval of the Director, Kenya Medical Research Institute. The project was funded by Gavi, The Vaccine Alliance. TNW (202800) and JAGS (098532) were funded by Wellcome Trust Fellowships and the KEMRI–Wellcome Trust Research Programme in Kilifi, Kenya received core support (203077) from the Wellcome Trust.

## Author contributions

JAGS and OSL conceived the study; JI, KeM, TK, ShS, MDK, OSL, LLH and JAGS designed the study; MS, JI, SK, SyS, VO, LLH and NM obtained and prepared the clinical and radiological data; MS, KaM, JI, JS, KP, RB and FG reviewed and categorised the radiographic images; AM, TB, IA, EB, MaO, TN and JAGS obtained and analysed demographic and vaccine clinic data; MiO, CB and JAGS undertook the time series analyses; MS and JAGS wrote the first draft of the manuscript; all authors critically reviewed the manuscript and approved the final draft.

---

**Suppl. table 1.** Incidence rate ratios for season, time and PCV10 introduction against hospitalisation with WHO-defined severe or very severe pneumonia among children aged 2-59 months.

| **Variable** | **Incidence Rate Ratio** | **95% CI** | **P value** |
| --- | --- | --- | --- |
| PCV10 introduction | 0·73 | 0·54, 0·97 | 0·033 |
| Time (in months) | 0·99 | 0·99, 1·00 | 0·001 |
| Seasonal month |  |  |  |
| January | Ref |  |  |
| February | 0·88 | 0·74, 1·05 | 0·150 |
| March | 0·73 | 0·58, 0·93 | 0·010 |
| April | 0·61 | 0·48, 0·77 | < 0·001 |
| May | 0·57 | 0·44, 0·73 | < 0·001 |
| June | 0·59 | 0·46, 0·76 | < 0·001 |
| July | 0·73 | 0·58, 0·93 | 0·009 |
| August | 0·73 | 0·57, 0·94 | 0·013 |
| September | 0·65 | 0·50, 0·84 | 0·001 |
| October | 0·74 | 0·55, 0·99 | 0·041 |
| November | 0·94 | 0·75, 1·18 | 0·605 |
| December | 1·09 | 0·91, 1·29 | 0·354 |

CI, confidence interval

**Suppl Table 2.** Correlation between annual admission counts of clinically-defined pneumonia and discharge diagnoses of other diseases in the pre-vaccine era (2002-2010)

| Diagnosis* | Number | Correlation | 95% CI | P-value |
| --- | --- | --- | --- | --- |
| ***At discharge*** |  |  |  |  |
| Unclassified | 274 | 0.85 | 0.42, 0.97 | 0.004 |
| Gastroenteritis | 2,146 | 0.81 | 0.32, 0.96 | 0.008 |
| Encephalopathy | 66 | 0.80 | 0.28, 0.96 | 0.010 |
| Trauma | 177 | 0.74 | 0.15, 0.94 | 0.023 |
| Malnutrition | 767 | 0.67 | 0.02, 0.92 | 0.046 |
| Febrile convulsions | 521 | 0.56 | -0.16, 0.89 | 0.11 |
| Meningitis | 67 | 0.52 | -0.22, 0.88 | 0.15 |
| Malaria | 2,308 | 0.48 | -0.27, 0.87 | 0.19 |
| Sepsis | 89 | 0.45 | -0.31, 0.86 | 0.23 |
| All other conditions | 632 | 0.40 | -0.36, 0.84 | 0.28 |
| Burns | 369 | 0.35 | -0.41, 0.82 | 0.36 |
| Anaemia | 399 | 0.32 | -0.44, 0.81 | 0.41 |
| Cellulitis | 280 | 0.22 | -0.52, 0.77 | 0.57 |
| Kerosene poisoning | 97 | 0.20 | -0.53, 0.76 | 0.60 |
| Upper Respiratory Infection | 322 | 0.16 | -0.56, 0.75 | 0.68 |
| Epilepsy | 121 | 0.08 | -0.62, 0.71 | 0.84 |
| Sickle cell disease | 243 | 0.02 | -0.65, 0.68 | 0.95 |
| Elective surgery | 132 | -0.11 | -0.72, 0.60 | 0.79 |
| ***On admission*** |  |  |  |  |
| Diarrhoea | 3,010 | 0.89 | 0.57, 0.98 | 0.001 |

CI confidence interval
- \* Between 2002-2010 there were 28,000 paediatric admissions in the age group 2-59 months; 16,245 (58%) were residents of the KHDSS. Of these 6,346 had a clinical diagnosis of severe or very severe pneumonia. Of the remaining 9,899 children, 643 had a primary discharge diagnosis of lower respiratory tract infection (LRTI) and 272 had a secondary discharge diagnosis of LRTI; the discharge diagnoses of 70 children were missing. All of these were excluded from the analysis leaving 8,914 children with non-pneumonia diagnoses. As gastroenteritis was the classified category with the highest correlation, we also examined the correlation between admission with diarrhoea (but without severe or very severe pneumonia) and admission with pneumonia which is given in the last row of the table.

**Suppl table 3.** Incidence rate ratios for the effects of PCV10 introduction against hospitalisation with WHO-defined severe or very severe pneumonia among children aged 2-143 months, in sub-groups of age and clinical severity, and with diarrhoea and all other admissions.

| End-point and sub-groups | PCV10 Incidence Rate Ratio | 95% CI | P value |
| --- | --- | --- | --- |
| <u>Severe pneumonia</u> |  |  |  |
| 2-59 months | 0·60 | 0·40, 0·91 | 0·017 |
| <u>Very severe pneumonia</u> |  |  |  |
| 2-59 months | 0·87 | 0·56, 1·34 | 0·519 |
| <u>Severe or very severe pneumonia</u> |  |  |  |
| 2-59 months | 0·73 | 0·54, 0·97 | 0·033 |
| 2-11 months | 0·70 | 0·50, 1·00 | 0·048 |
| 12-23 months | 0·84 | 0·61, 1·15 | 0·283 |
| 24-59 months | 0·71 | 0·43, 1·19 | 0·192 |
| 60-143 months | 0·95 | 0·56, 1·59 | 0·832 |
| <u>Diarrhoea</u> |  |  |  |
| 2-59 months | 0·63 | 0·31, 1·26 | 0·190 |
| 2-11 months | 0·79 | 0·42, 1·48 | 0·459 |
| 12-23 months | 0·63 | 0·41, 0·99 | 0·044 |
| 24-59 months | 0·62 | 0·39, 0·99 | 0·046 |
| 60-143 months | 0·66 | 0·46, 0·95 | 0·027 |
| <u>All other admissions</u> |  |  |  |
| 2-59 months | 0·89 | 0·57, 1·39 | 0·614 |

CI, confidence interval

